# SMART: A Somatic Mutation Annotation and Reporting Tool for cancer genomics

**DOI:** 10.64898/2026.07.15.738659

**Authors:** M. Dominguez, I. Reddin, J. Gibson, M. Rudraraju, K. Veal, C. Kipps, A.P. Williams, S Ennis

**Author notes:** Corresponding authors: Dominguez, M. & Reddin, I. Manuel -; Ian.

## Abstract

**Motivation:** Translational interpretation of somatic variants from targeted oncology panels is hampered by inconsistent transcript prioritisation and by the need for reproducible pipelines that natively integrate OncoKB-derived evidence for research purposes.

**Results:** We present SMART (Somatic Mutation Annotation and Reporting Tool), a Dockerised pipeline that embeds OncoKB API-derived annotations, including therapeutic (L1–4), resistance (R1–R3), diagnostic (Dx1–3), prognostic (Px1–3) and FDA levels, directly into a VCF-based workflow. SMART combines this with VEP, CIViC, Cancer Hotspots, ClinVar, SpliceAI, REVEL, LOEUF and gnomAD, applies a unified three-tier transcript prioritisation (whitelist > MANE Select > VEP fallback), and produces three-tiered outputs for computational, bioinformatic and research interpretation. Validation against reference APIs showed full concordance across 804 field-level checks.

**Availability and Implementation:** Source code and Docker image are freely available at https://github.com/WeTGI-colab/SMART under the MIT License. SMART is provided for research use only; use of SMART outputs for patient specific clinical reports, clinical decision-making, or other patient-facing purposes requires appropriate governance and all required third-party licensing, including any OncoKB licence required for patient report generation.

## 1 Introduction

The integration of next-generation sequencing into routine oncology practice has created a pressing need for robust, reproducible frameworks that can translate raw variant calls into clinically meaningful information. Targeted oncology sequencing panels (e.g. TruSight Oncology 500 v2, Oncomine Precision Assay, RMH200 Solid Tumour DNA ) [1], are increasingly used in research and translational settings to interrogate predefined sets of cancer-relevant genes, generating somatic variant calls that require systematic functional and curated-evidence interpretation before they can support research analysis and hypothesis generation. A range of tools has been developed for the annotation of somatic genomic variants, including general-purpose annotators such as the Ensembl Variant Effect Predictor (VEP) [2] and ANNOVAR [3], as well as cancer-focused tools and workflows such as Oncotator [4], CancerVar [5], and PCGR [6]. These tools address important parts of the annotation workflow, but typically emphasise different objectives, including consequence annotation, transcript- normalised conversion to Mutation Annotation Format (MAF), or comprehensive multi- layer cancer annotation and research summarisation that combines variant annotation with genome-level summary metrics such as tumour mutational burden, microsatellite instability status, and mutational signatures.

In parallel, several specialised knowledge bases and predictive resources have emerged to support interpretation of somatic variants by curating functional, biological, and curated evidence from the literature and large-scale cancer datasets. These include the Clinical Interpretations of Variants in Cancer (CIViC) [7], OncoKB [8], Catalogue of Somatic Mutations in Cancer (COSMIC) [9], Cancer Hotspots [10], and ClinVar [11], alongside in silico predictors and constraint metrics such as SpliceAI [12], REVEL [13], and loss-of-function observed/expected upper bound fraction (LOUEF) [14]. Collectively, these resources provide complementary evidence relating to tumorigenic relevance, therapeutic relevance, splicing impact, predicted pathogenicity, and gene-level intolerance. When integrated in a consistent manner, they can substantially improve the interpretability of somatic variant calls.

Despite these advances, several challenges remain when applying existing annotation pipelines in research and translational settings. First, outputs are often optimised either for computational analysis or for research-oriented review, rather than for research use by bioinformaticians, molecular scientists and translational research teams. Second, transcript selection is frequently a source of inconsistency in somatic variant reporting, with different tools and databases prioritising different transcripts or consequences for the same genomic event. Third, clinically-relevant curated annotations are often distributed across multiple resources that vary in curation practice, update frequency, and access model, requiring additional downstream harmonisation before results can be reviewed in a research-interpretable format. Finally, local deployment, version control, and reproducibility are essential requirements in many clinical and research genomics environments, where reliance on fragmented multi-tool workflows can complicate validation, governance, and reproducible implementation.

To address these challenges, we developed SMART (Somatic Mutation Annotation and Reporting Tool), a fully Dockerised pipeline for the systematic and structured research annotation of somatic variants identified from targeted sequencing panels. SMART integrates consequence annotation via VEP with somatic-focused resources, including CIViC, Cancer Hotspots, COSMIC, ClinVar, SpliceAI, REVEL, and LOEUF, and augments these with curated oncogenicity classifications and therapeutic associations retrieved from OncoKB. A unified transcript prioritisation strategy is applied throughout the workflow to ensure consistency between annotation, downstream interpretation, and report generation. SMART produces three structured outputs tailored to distinct end users: a comprehensive MAF-formatted file for computational analysis, a detailed annotation table for bioinformatic review, and a clinical research-focused subset intended to support variant review in translational and multidisciplinary research settings. Although SMART was developed and validated using VCF files generated from TSO500 data processed with the DRAGEN variant calling pipeline, it is designed to accept standard-compliant VCF input and is therefore intended for broader use across targeted sequencing assays and variant callers. By packaging all components within a standardised Docker container and providing auxiliary scripts for reference data management, SMART enables reproducible, high-throughput somatic variant annotation within local research genomics workflows. Here, we describe the design, implementation, and validation of SMART, and assess its performance as a reproducible framework for somatic variant annotation and reporting in targeted-panel oncology workflows.

## 2 System and methods

### 2.1 Pipeline overview

SMART is organised as a three-step workflow (Figure 1): (i) variant consequence annotation using the VEP, extended through plugins to include splice impact (SpliceAI), missense pathogenicity (REVEL), gene-level loss-of-function intolerance (LOEUF), clinical assertions (ClinVar), and somatic-focused resources (CIViC, Cancer Hotspots, gnomAD); (ii) curated evidence annotation using OncoKB to retrieve oncogenicity classifications, mutation effects, and curated evidence levels; and (iii) output generation, in which annotated variants are consolidated into three structured outputs aimed at distinct end users: a complete MAF for downstream cohort-level and programmatic analysis, a detailed annotation table for interactive bioinformatic review, and a focused subset for research interpretation in multidisciplinary research settings.

**Figure 1.**
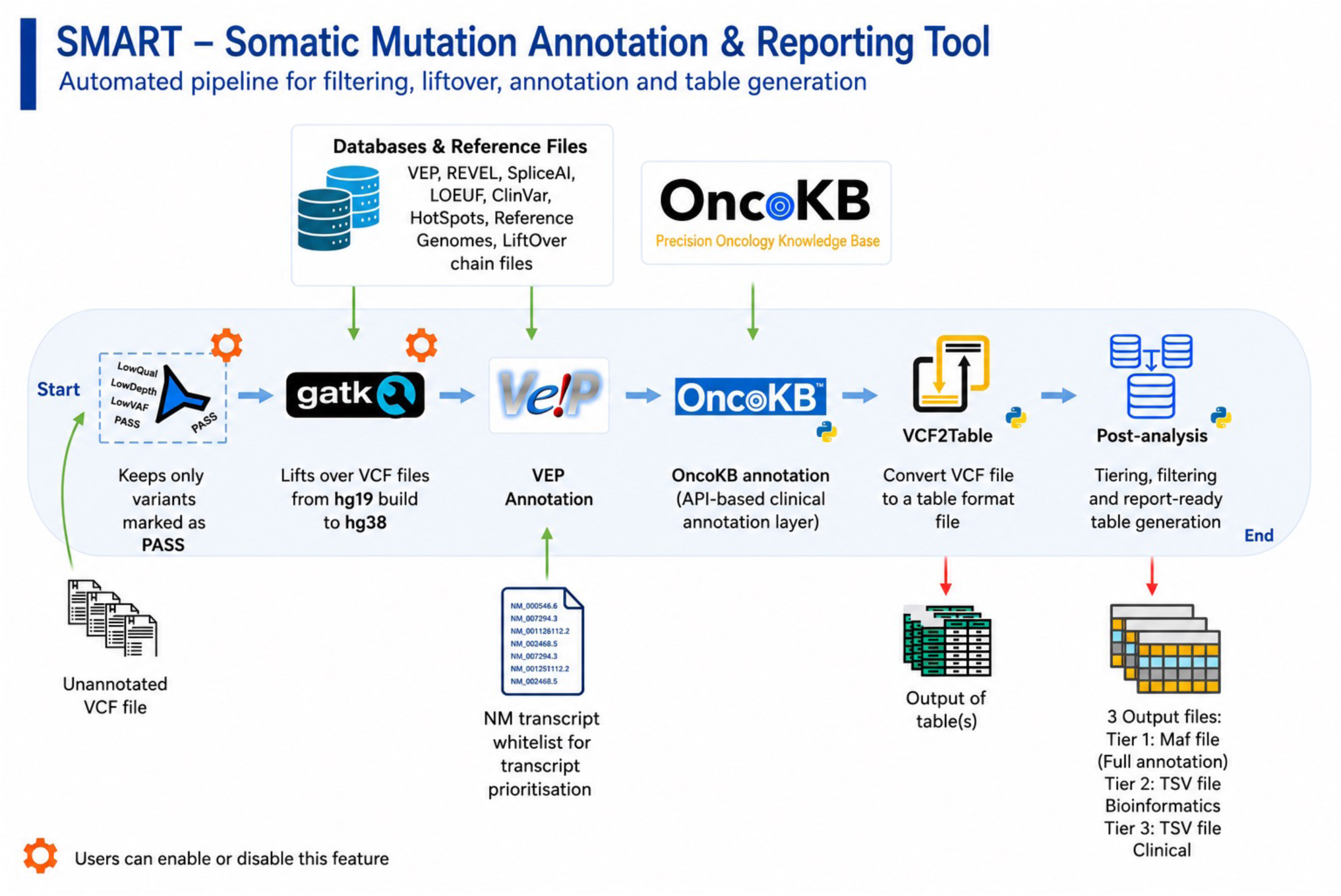
Overview of the SMART workflow. SMART (Somatic Mutation Annotation & Research Tool) is an automated pipeline for somatic variant filtering, liftover, annotation, and analysis-ready table generation. Starting from an unannotated VCF file, SMART retains only PASS variants and optionally performs hg19-to-hg38 liftover using GATK. Variants are subsequently annotated with Ensembl VEP using multiple reference databases and plugins, including ClinVar, SpliceAI, REVEL, LOEUF, Cancer Hotspots, and reference genome resources. A transcript-prioritisation layer based on a user-defined NM transcript whitelist is incorporated during annotation. SMART then performs API-based curated-evidence annotation through OncoKB before converting the annotated VCF into tabular format. Post-analysis modules perform filtering, tiering, and generation of analysis-ready outputs. The pipeline produces three principal outputs: (i) a complete MAF annotation file (Tier 1), (ii) a TSV file for bioinformatics review (Tier 2), and (iii) a research-oriented TSV evidence summary (Tier 3). Orange gear icons indicate optional workflow components that can be enabled or disabled by the user.

All software components are integrated and distributed within a single Docker container, ensuring that all dependencies are version-locked and packaged within a standardised, reproducible execution environment. The application includes auxiliary scripts to automate the download and configuration of the required reference datasets, described in Section 4.

### 2.2 Input requirements and coordinate handling

SMART accepts one or more single-sample VCF files encoding the genomic coordinates of somatic single-nucleotide variants (SNVs), small insertions/deletions (indels), and structural variants. Only variants flagged as PASS in the FILTER column are retained for downstream processing; all other entries are discarded before annotation, unless filtering is disabled.

The pipeline natively supports the human reference genome assembly GRCh38. To accommodate legacy GRCh37 inputs, an optional pre-processing step performs hg19- to-hg38 coordinate conversion using GATK LiftoverVcf (v4.6) and the UCSC chain file. This liftover step is configurable and can be enabled or disabled by the user.

### 2.3 Variant consequence annotation

Variant consequence annotation is performed using Ensembl VEP, configured with a fixed set of parameters to ensure consistency and reproducibility across runs. The configuration includes gene cross-references, protein domain annotations, and overlap with regulatory regions, using GENCODE as the underlying transcript model. All transcript-dependent consequences for each variant are retained in the VEP-annotated VCF output; downstream selection of the transcript to be displayed in downstream outputs is governed by the prioritisation strategy described in Section 3 (Algorithm).

### 2.4 Annotation resources

In addition to the VEP-derived consequence layer, SMART integrates a curated set of complementary annotation resources to support research interpretation. ClinVar provides germline and somatic clinical assertions; CIViC provides variant-level and evidence-level cancer interpretations; COSMIC provides curated somatic cancer mutation annotations and was used to flag variants previously reported in tumour samples; Cancer Hotspots provides linear and three-dimensional hotspot annotations; SpliceAI provides splice-altering predictions; REVEL provides ensemble-based missense pathogenicity scoring; LOEUF (gnomAD v4.0) provides gene-level loss-of-function intolerance; gnomAD exome and genome cohorts provide population allele frequencies; and OncoKB provides curated oncogenic classifications, mutation effects, and the full hierarchy of curated evidence levels (therapeutic sensitivity L1–4, resistance R1–R3, diagnostic Dx1–3, prognostic Px1–3, and FDA regulatory levels).

ClinVar, CIViC, COSMIC, Cancer Hotspots, SpliceAI, REVEL, LOEUF, and gnomAD are accessed locally via VEP plugins and pre-staged reference files. OncoKB is queried programmatically via its REST API; the integration mechanism is described in Section 4.1. The Docker container does not bundle the OncoKB token, which must be obtained by the user and supplied at runtime in accordance with OncoKB licensing terms. The reference installer script is described in Section 4.2.

## 3 Algorithm

### 3.1 Unified transcript prioritisation

A central design choice of SMART is consistent transcript prioritisation across all annotation and reporting steps. Because somatic variants frequently map to multiple overlapping transcripts with divergent protein-level consequences, the selection of which transcript to query against curated knowledgebases (and which to display in the final output) has direct implications for the output oncogenicity classification, therapeutic evidence level, and prognostic associations assigned to a given variant (demonstrated empirically in Section 4.5.2). To remove this source of inconsistency, SMART applies a single three-tier prioritisation scheme at both the OncoKB annotation step (Section 4.1) and the table-generation step (Section 4.3).

For each variant, transcripts reported by VEP are evaluated in the order below; the first tier that returns a match determines the transcript used downstream.

**Tier 1 — User-provided whitelist.** RefSeq NM accessions present in the user-supplied transcript list. Matching is version-agnostic (e.g. NM_005228 matches NM_005228.5), so that panel-specific or laboratory-curated preferred transcripts continue to apply across RefSeq releases without requiring manual revision. The whitelist is supplied as a plain-text file at runtime.

**Tier 2 — MANE Select / MANE Plus Clinical.** If no Tier 1 match exists, the transcript annotated by Ensembl/NCBI as MANE Select or MANE Plus Clinical is selected. This guarantees that, in the absence of a panel-specific preference, the variant is reported on the transcript jointly recommended by Ensembl and NCBI for consistent variant interpretation.

**Tier 3 — VEP fallback.** If neither Tier 1 nor Tier 2 criteria are satisfied, the first transcript reported by VEP for that variant is retained, preserving compatibility with the default behaviour of VEP for variants outside curated transcript sets. In practice, because MANE Select and MANE Plus Clinical transcripts are already exhausted by Tier 2, Tier 3 resolution is most often determined by canonical status, APPRIS annotation, transcript support level, biotype, consequence severity rank, and transcript length.

The selected transcript governs both (i) the HGVS protein notation submitted to the OncoKB API and (ii) the transcript-specific fields populated in the Tier 2 and Tier 3 outputs. This guarantees that the transcript shown to the reviewer is the same transcript on which the curated annotation was retrieved, eliminating a class of inconsistency that is otherwise undetectable in downstream review.

## 4 Implementation

### 4.1 OncoKB API integration and VCF reintegration

Although OncoKB annotations are publicly accessible, automated large-scale integration requires the use of the OncoKB REST API, as the framework does not natively support VCF-formatted inputs. SMART addresses this limitation through an intermediate transformation layer that extracts variant-level features from the VEP-annotated VCF and reformats them into the per-variant input expected by the OncoKB API.

For each variant, the prioritised HGVS.p notation (Section 3.1) and gene identifier are submitted to the API to retrieve curated annotations, including oncogenicity classification, biological mutation effect, the full hierarchy of curated evidence levels (therapeutic sensitivity L1–4, resistance R1–R3, diagnostic Dx1–3, prognostic Px1–3, FDA regulatory), and tumour-type–specific drug associations. API responses are returned in structured JSON, parsed, and normalised into a flat set of fields that are reintegrated into the original VCF as additional INFO entries. The resulting VCF therefore consolidates VEP consequence-level annotations and OncoKB curated knowledge within a single, standard-compliant representation, preserving compatibility with all downstream tools that accept VCF input. To our knowledge, this direct integration of API-derived OncoKB annotations into a VCF-based annotation framework has not been previously described in the literature.

Variants are routed to the appropriate OncoKB endpoint based on type. SNVs, indels, MantaINS (structural insertions) and MantaBND (translocation breakends) are submitted to /annotate/mutations/byProteinChange, as these alterations produce specific protein-level consequences for which OncoKB curates distinct therapeutic associations (e.g. EGFR exon 20 insertions). Copy-number alterations (MantaDUP, MantaDEL, GAIN, LOSS) are submitted to /annotate/copyNumberAlterations, which accepts gene-level rather than residue-level inputs. For Copy Number Alterations (CNAs), the same three-tier transcript prioritisation scheme (Section 3.1) is applied to resolve the correct gene symbol when a single copy-number segment overlaps multiple transcripts, for example distinguishing *CDKN2A* from the adjacent *CDKN2B* in 9p21 deletions, before the gene identifier is submitted to OncoKB.

A known limitation of the public OncoKB MafAnnotator client is that it does not propagate CNA-level annotations into the MAF correctly. To address this, the post-analysis step of SMART overwrites the MafAnnotator-derived fields for CNA rows (VARIANT_IN_ONCOKB, ONCOGENIC, MUTATION_EFFECT, all therapeutic level columns, and HIGHEST_SENSITIVE_LEVEL / HIGHEST_RESISTANCE_LEVEL) with values returned directly by the OncoKB REST API, ensuring consistent annotation across SNVs, indels, structural variants and copy-number alterations within a single output table.

The OncoKB API requires a user-specific access token; SMART does not bundle a token in the distributed Docker image. Users must obtain a token in accordance with OncoKB licensing terms (https://www.oncokb.org/account/register) and supply it at runtime through the configuration file. Use of OncoKB-derived annotations must remain within the user’s permitted licence scope.

### 4.2 Docker container and reference data installer

All software components (VEP and its plugins, GATK, the OncoKB client, the table-generation scripts, and their dependencies) are distributed within a single Docker container. Dependencies are version-locked at image build time, providing a reproducible execution environment and removing the need for users to manage individual tool installations. The container is published with a fixed tag corresponding to the manuscript version (smart:vX.Y) and is accompanied by a configuration file (Config.yaml) that exposes all runtime parameters, including reference paths, the transcript whitelist, OncoKB token, and the per-tier annotation field selection used to compose the output files.

Reference data setup is managed by a dedicated installer script (utils/get_ref_files.sh), which downloads and prepares all required annotation resources in a single command: the GRCh38 reference genome and UCSC liftover chain, ClinVar, CIViC, REVEL, gnomAD constraint metrics (LOEUF), Cancer Hotspots (linear and 3-D), and the Ensembl VEP cache and associated plugins. The installer is idempotent, meaning completed steps are detected and skipped on subsequent runs and records database versions and download dates in an auto-generated configuration file, ensuring that the exact reference state used for any analysis can be reconstructed. SpliceAI scores, which require an Illumina BaseSpace account, must be staged manually; the installer detects their absence and emits explicit download instructions.

### 4.3 Output tiers

SMART produces three structured output files at the end of each pipeline run, each targeting a distinct analytical use case. The composition of Tier 2 and Tier 3 is fully configurable through Config.yaml, in which every annotation field is assigned an explicit tier designation. Both subsampled files include a two-row header: the first row contains field names; the second row provides human-readable metadata for each field, comprising a plain-language description, the originating data source, and the version or release identifier. This design enables downstream users to interpret annotation content without cross-referencing external documentation, and supports auditability by embedding provenance information directly within the output file.

Tier 1 is a complete MAF file retaining all 1,028 annotation fields generated by SMART. It encompasses the full set of VEP consequence fields, all OncoKB annotation fields, ClinVar, CIViC, Cancer Hotspots, SpliceAI delta scores, REVEL, LOEUF, gnomAD population frequencies, MAF-standard columns (Hugo_Symbol, Variant_Classification, HGVSp_Short, and related identifiers), and all per-sample FORMAT fields carried over from the input VCF. Tier 1 is intended for use in computational downstream analyses, including cohort-level somatic mutation profiling, integration with MAF-based analytical frameworks, and custom filtering or aggregation workflows. A breakdown of the 1,028 fields by annotation source and category is provided in Supplementary Figure S1.

Tier 2 retains 670 fields in a tab-delimited format and is designed for detailed bioinformatic review. It preserves relevant annotation content while excluding redundant MAF scaffolding columns, providing a technically complete view of all variant-level annotations alongside their embedded metadata. Tier 2 is the primary output for our research bioinformaticians performing annotation quality checks, troubleshooting transcript assignments, or extracting specific annotation layers for downstream processing.

Tier 3 is a 77-field research-focused subset presented in the same TSV format with the two-row annotated header. Fields are selected to support research variant review in our multidisciplinary research team, without requiring familiarity with the full annotation space. The subset includes: core variant identity fields (sample identifier, genomic coordinates, reference and alternate alleles, HGVS coding and protein notation, prioritised transcript); VEP-derived consequence and impact tier; in silico pathogenicity and splicing predictors (SpliceAI, REVEL, LOEUF, SIFT, PolyPhen); population allele frequencies from gnomAD exome and genome cohorts; ClinVar clinical significance and oncogenicity assertions; CIViC variant-level and evidence-level fields; Cancer Hotspots linear and three-dimensional annotation; COSMIC curated somatic cancer mutation annotations for variants previously reported in tumour samples; OncoKB oncogenicity classification and mutation effect; and the full hierarchy of OncoKB evidence levels, including therapeutic sensitivity (Level 1–4), therapeutic resistance (R1–R3), diagnostic (Dx1–3), prognostic (Px1–3), and FDA regulatory levels. Variant allele fraction (VAF) is also included to contextualise tumour burden. Table 1 provides a schematic summary of the SMART output tiers and the annotation fields retained in the current configuration of each output format.

**Table 1.**
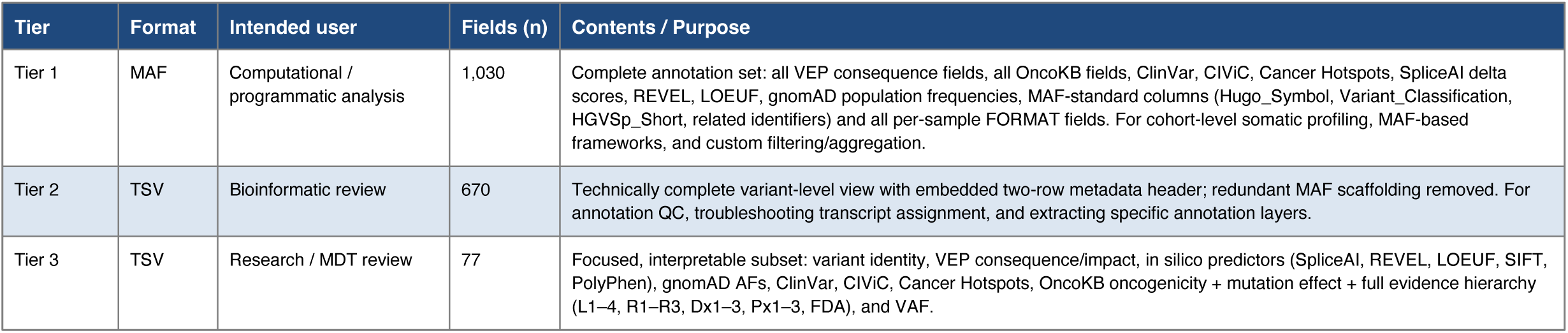
Summary of SMART output tiers. Fields (n) denotes the number of annotation columns retained in each output file. MAF, Mutation Annotation Format; TSV, tab-separated values; VEP, Variant Effect Predictor; MDT, multidisciplinary team; VAF, variant allele fraction.

### 4.4 Auxiliary outputs and run logging

To support quality control and reproducibility, SMART generates two auxiliary outputs alongside the main annotation results. A variant count summary file records, for each sample, the number of variants present at every stage of the pipeline: original input, PASS filtering, liftover, VEP annotation, OncoKB annotation, and final table generation to enabling rapid identification of any step at which variants are unexpectedly lost. A run log captures the SMART version, the version of every reference resource and tool used, step-level progress messages, and the configuration parameters of the run. Together, these outputs facilitate routine quality monitoring and retrospective audit, which are particularly relevant when SMART is deployed in clinical genomics settings where traceability is essential.

### 4.5 Validation

To assess the analytical fidelity, robustness, and computational characteristics of SMART, three independent verification sets were designed: field-level annotation concordance against reference APIs (4.5.1), the impact of transcript prioritisation on OncoKB-derived evidence annotation (4.5.2), compatibility across heterogeneous somatic variant callers (4.5.3), and runtime characterisation on standard hardware (4.5.4). The complete verification suite is provided in the SMART repository and can be re-executed end-to-end by any user.

#### 4.5.1 Annotation concordance against reference APIs

Annotation fidelity was assessed by comparing SMART outputs against the corresponding reference APIs for three independent modules: OncoKB (v7.0), Ensembl VEP (v114.0), and the Clinical Interpretation of Variants in Cancer (CIViC) database. The curated input panel comprised 18 variants (14 SNVs and indels covering oncogenic hotspots across canonical cancer genes, and 4 copy-number alterations) which generated 22 output rows due to multiple preferred-isoform matches in a subset of variants (Section 3.1; gene-level composition in Tier 1). Across the three modules, 804 field-level checks were performed, with complete concordance and no coverage gaps relative to the reference APIs (Table 2). The corresponding test scripts are bundled with the SMART repository so that the same checks can be re-run by future users. Full variant-level and field-level results for all three modules are provided in <u>Supplementary</u> <u>Data 1</u>.

**Table 2.** Field-level concordance of SMART pipeline annotations against reference APIs. Output rows assessed denote the number of output rows evaluable for a given annotation source; the 18 input variants in the curated panel produced 22 output rows due to multi- transcript expansion for variants overlapping multiple whitelist isoforms.

| Annotation source | Output rows assessed | Fields per variant | Field checks (n) | Concordant checks | Notes |
| --- | --- | --- | --- | --- | --- |
| OncoKB | 22 | 23 | 506 | 506 (100%) | 3 fields untestable (tumour-type-dependent) |
| VEP | 17 | 14 | 238 | 238 (100%) | 5 variants excluded: 4 CNAs, 1 missing HGVSc |
| CIViC | 18 | 4 | 56 | 56 (100%) | 4 CNAs excluded; |
| Total | 22 | — | 804 | 804 (100%) | 0 mismatches |

#### 4.5.2 Impact of transcript prioritisation on OncoKB-derived evidence

To demonstrate that transcript selection has direct and reproducible consequences on OncoKB-derived evidence annotation, the pipeline was executed twice on an identical input VCF using two distinct transcript configurations: a set of panel-curated preferred transcripts (set A) and a set of biologically plausible but non-preferred alternatives (set B). Across four representative variants, the choice of transcript determined protein consequence, OncoKB oncogenicity classification, and the presence or absence of therapeutic and prognostic evidence levels (Table 3). Because downstream knowledgebase queries are driven by the protein change or gene identifier submitted, use of alternative transcripts caused variants with OncoKB therapeutic evidence to be presented to OncoKB under unrecognised alterations or under the wrong gene entirely, abrogating all therapeutic and diagnostic evidence annotations.

**Table 3.**
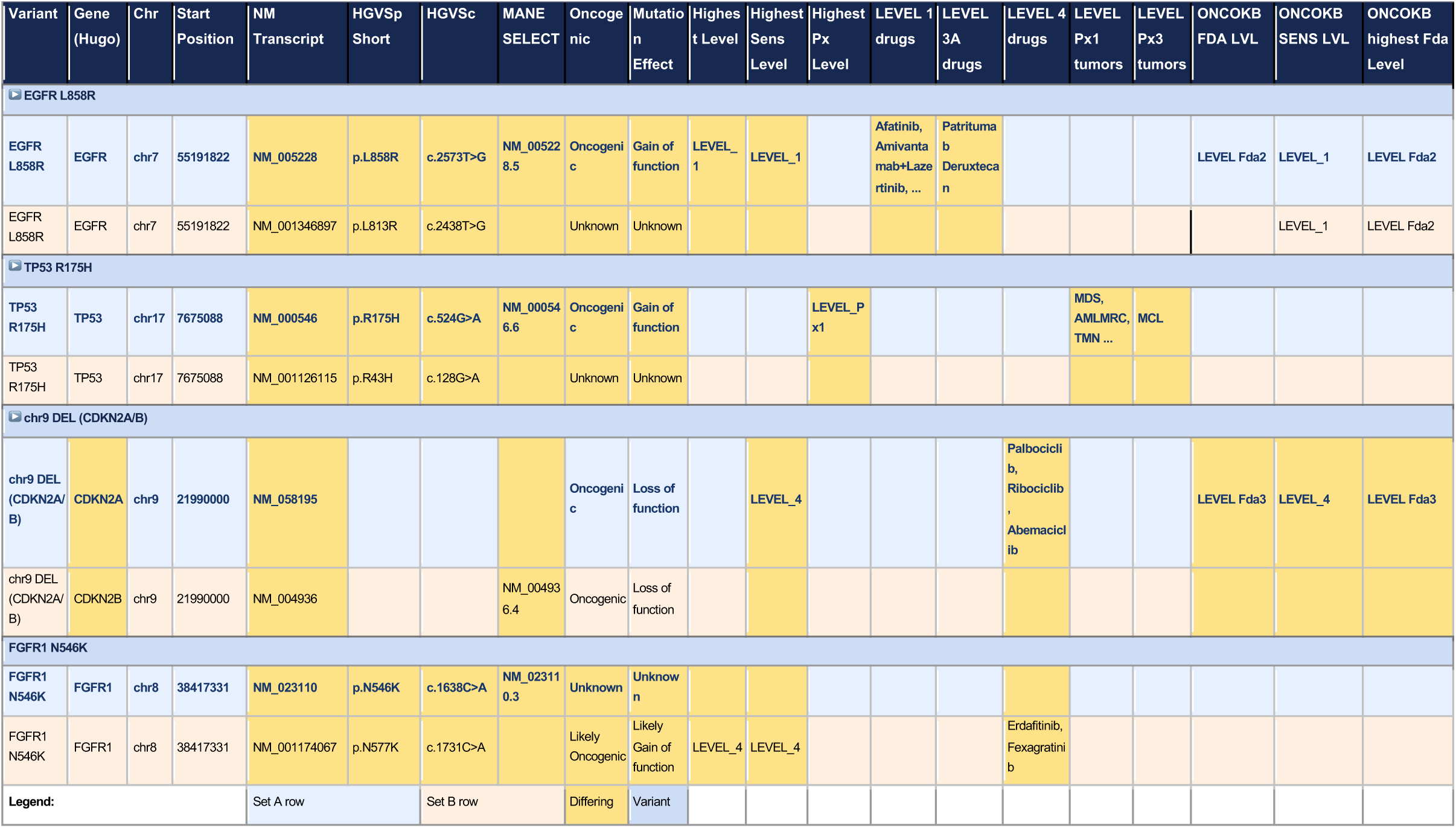
Transcript selection determines OncoKB-derived evidence annotation across four representative variants. The SMART pipeline was run twice on the same VCF using two different transcript whitelists (set A, panel-curated preferred; set B, alternative isoform), and the resulting OncoKB annotations compared.

These four cases illustrate that transcript choice can drive OncoKB-derived evidence annotation in both directions. The *EGFR* L858R example is a direct false negative: the canonical NM_005228.5 transcript yields p.Leu858Arg with LEVEL_1 actionability and multiple FDA-approved agents, whereas the alternative NM_001346897.2 places the same genomic substitution at codon 813 (p.Leu813Arg), an alteration absent from OncoKB and therefore returned as *Unknown*, silently losing all therapeutic evidence. The *FGFR1* N546K example is the converse: the TSO500 panel mandates a non-MANE isoform (NM_001174067.2) under which the variant becomes p.Asn577Lys, returned by OncoKB as Likely Oncogenic with LEVEL_4 evidence, while the MANE Select transcript (NM_023110.3) yields p.Asn546Lys, currently unclassified. The therapeutic signal, therefore, exists or does not exist depending entirely on isoform selection, with neither outcome flagged as uncertain. Tools that enforce a fixed transcript hierarchy without panel-specific overrides (e.g. PCGR, which does not permit user-specified preferred transcripts) cannot guarantee that the reported annotation aligns with either the panel design or the assumptions underpinning the knowledgebase entries. The use of additional transcripts beyond MANE transcripts is not only supported by best-practice recommendations for bioinformatics approaches applied to high-throughput sequencing [15], but is also common practice in genomics workflows.

#### 4.5.3 Compatibility across variant callers

The primary development focus of SMART was the processing of DRAGEN-derived somatic VCF files. To ensure that the tool not only met the needs of our team but also those of future users, a verification strategy was implemented.

Variant caller compatibility was assessed using four VCF files from the SEQC2 benchmark dataset (tumour/normal pair FFG_GZ_T_24h-B versus WGS_IL_N_1), generated using GATK MuTect2, Illumina Strelka, SomaticSniper, and the tumour-only amplicon caller Pisces (v5.2.10). Each VCF was processed through SMART to confirm structural output integrity, correct handling of PASS filters, and consistent VEP and OncoKB annotations across callers with differing FORMAT field conventions.

Variant caller–specific annotation fields not defined in the configuration file were correctly propagated into the three-tiered output files, with their descriptions flagged as “UNKNOWN”. This is expected. Future users with annotation-specific VCF files need to modify the configuration file accordingly.

#### 4.5.4 Runtime and scaling

To improve scalability, SMART was created with support for parallel processing through the -jobs parameter, enabling the simultaneous analysis of multiple VCF files. Benchmarking was performed using 100 VCF files comprising more than 250,000 variants in total on a desktop workstation equipped with an Intel Core i7-3770S processor (8 cores, 3.10 GHz) and 15 GB RAM. Using 8 parallel jobs, the complete analysis was completed in 178 minutes.

As part of our effort to develop a robust and portable tool capable of operating across a wide range of computing environments, memory consumption was also monitored throughout the benchmark. The results demonstrated a modest memory footprint, with peak memory usage remaining below 11 GB during the analysis (Supplementary Figure S2).

### 4.6 Reproducibility and portability

The verification suite described in Section 4.5 has been executed end-to-end across multiple computational environments, including a desktop workstation (Intel Core i7- 3770S, 8 threads, 3.10 GHz, 15 GB RAM), a personal laptop (MacBook Air, Apple M5, 16 GB RAM), and the IRIDIS6 high-performance computing cluster at the University of Southampton. Identical results were obtained on all platforms, confirming that the Docker-based deployment is portable across heterogeneous hardware. Together with the version-controlled installer (Section 4.2), this enables any user to deploy a fully reproducible instance of SMART from a single repository, archived at Zenodo (DOI: 10.5281/zenodo.20445865).

### 4.7 Comparison with existing pipelines

Several frameworks have been developed to support somatic variant interpretation in cancer genomics, of which PCGR and CancerVar are among the most widely adopted. To position SMART relative to these tools, a structured comparison was performed across functional annotation, in silico prediction, population and somatic databases, curated interpretation frameworks, supported variant types, and technical implementation (Supplementary Figure S3). All three tools share VCF-based input and produce annotations for SNVs and indels but differ substantially in their breadth of curated interpretation and the configurability of upstream choices. The most consequential differentiator is SMART’s native integration of the full OncoKB evidence hierarchy which is not natively available in either PCGR or CancerVar. Both alternatives rely instead on evidence aggregation from CIViC/CGI or on internal rule-based scoring (OPAI in CancerVar), neither of which produces the same standardised therapeutic tier assignments used in published evidence hierarchies. SMART is also unique among the three in supporting panel-specific transcript whitelists, built-in hg19→hg38 liftover, and audience-targeted tiered outputs (MAF/TSV) with embedded provenance metadata. In contrast, PCGR emphasises broad annotation and evidence aggregation, and CancerVar focuses on automated guideline-based scoring; SMART prioritises locally deployable, research-focused, evidence-rich interpretation through curated knowledgebase integration.

## 5 Discussion

SMART addresses a practical gap between raw somatic variant annotation and the structured outputs needed in routine research genomics workflows. Existing frameworks provide comprehensive functional and curated evidence but typically optimise their output for downstream computational analysis or research-oriented exploration, rather than for structured review by multidisciplinary research teams. SMART bridges this gap by consolidating consequence annotation, somatic-focused resources, and OncoKB- curated evidence into a single workflow with tiered, audience-targeted outputs.

A central design feature is the unified transcript prioritisation strategy applied identically at the OncoKB annotation step (Section 4.1) and the table-generation step (Section 4.3). As demonstrated by the *EGFR* L858R and *FGFR1* N546K cases (Section 4.5.2), transcript choice can determine whether a variant with therapeutic evidence is recognised or silently lost, and inconsistency between the transcript queried and the transcript displayed is otherwise undetectable in research review. Enforcing a single prioritisation logic across the pipeline removes this class of error.

In current practice, molecular scientists and bioinformaticians frequently complement automated annotation by manually cross-checking variants of interest across multiple online resources, for example looking up individual alterations in the OncoKB, ClinVar, CIViC, COSMIC, and gnomAD web interfaces, reconciling differences in transcript representation and nomenclature, and recording the consulted database versions for traceability. This process is time-consuming, error-prone, and difficult to standardise across reviewers, particularly when cohorts contain large numbers of variants or when multiple analysts contribute to the same research analysis. By integrating VEP, CIViC, Cancer Hotspots, ClinVar, SpliceAI, REVEL, LOEUF, gnomAD, and OncoKB within a single workflow, and by embedding the source and version of each annotation directly in the output as part of the two-row header, SMART substantially reduces the need for such manual lookup and harmonisation. Reviewers can interrogate the relevant evidence for each variant in a single tabular view, with explicit provenance for every annotation field, which improves both efficiency and consistency in somatic variant annotation.

The Dockerised, version-controlled architecture supports reproducibility and simplifies deployment in local research genomics environments where governance, validation, and auditability are required (for example, for auditable bioinformatic tool version control). Combined with the reference-installer script, the variant-count summary, and the run log, these elements enable the exact analytical state of any past run to be reconstructed. Demonstrated portability across a desktop workstation, a personal laptop, and a high-performance computing cluster (Section 4.6) confirms that SMART can be deployed without specialised infrastructure.

SMART is particularly well-suited to targeted-panel workflows where the analytical goal is not only broad annotation but also the generation of concise, interpretable outputs for multidisciplinary research discussion. The three-tier output design (full MAF for computation, detailed TSV for bioinformatic review, and a research-focused TSV for interpretation) allows the same pipeline to serve distinct end users without separate downstream processing. The 100% concordance across 804 field-level checks against OncoKB, VEP and CIViC reference APIs (Section 4.5.1) and successful processing of VCFs from four heterogeneous somatic callers (Section 4.5.3) together support SMART as a reliable, caller-agnostic framework for routine somatic reporting in targeted-panel oncology.

## 6 Limitations

SMART has a substantial reference-data footprint, with the combined VEP cache, plugin resources, and ancillary databases requiring more than 100 GB of local storage, which may constrain deployment in storage-limited environments. Runtime and computational demands also scale with batch size and the breadth of the annotation configuration; SMART mitigates this through cohort-level batch processing with parallel per-sample execution and through configurable activation of the liftover and OncoKB modules, but very large cohorts will still benefit from deployment on shared computing resources.

Several of the integrated resources are knowledgebase-dependent and evolve over time, so identical input data analysed against different releases of OncoKB, ClinVar, CIViC, or VEP may yield different annotations. Components that depend on OncoKB additionally require appropriate access credentials and licensing: although OncoKB is freely accessible for academic research and provides a credentialed academic API, clinical and commercial use of the data, including annotation of patient sequencing reports, requires a paid licence (OncoKB Licensing FAQ). Users should also note that the FDA partial recognition of OncoKB granted in 2021 explicitly excludes variant-to-therapy mapping and is not intended to support reimbursement decisions. More broadly, the utility of any annotation pipeline depends on the completeness and curation quality of its underlying databases, which vary between genes, variant classes, and tumour types. These licensing constraints are a principal reason that SMART is framed here as a research tool rather than a clinical reporting system.

SMART itself is distributed under MIT licence and consists of code only; reference datasets are not redistributed within the container but are downloaded at install time via the bundled installer script. Most integrated resources (including VEP, GATK, CIViC, ClinVar, gnomAD/LOEUF, MANE, and GENCODE) are released under permissive licences or in the public domain, allowing free use in research settings and in other settings subject to their individual licences. Four resources require additional attention for clinical, patient-facing, or commercial deployment: OncoKB, which requires a paid licence for clinical or patient-facing use; SpliceAI, whose trained models and precomputed scores are released under CC BY-NC 4.0 and therefore require a commercial licence from Illumina for any non-academic use; and REVEL, whose scores are free for non-commercial use only and require direct permission from the developers for commercial application: and COSMIC, whose data are free for academic and non-commercial research use under user registration but require a commercial licence for commercial or clinical use. Because COSMIC data are not redistributable, they are neither bundled in the container nor retrieved by the installer; users must download COSMIC directly under their own registration and generate the reference file locally using the provided script. Users considering any clinical or commercial use of SMART should obtain the appropriate licences and complete local validation and governance before issuing patient-facing reports.

Although SMART accepts standard-compliant VCFs and has been verified against MuTect2, Strelka, SomaticSniper, and Pisces outputs, development and validation focused primarily on TSO500/DRAGEN data, and broader evaluation across additional assays would further support generalisability. The current implementation is best suited to SNVs, indels, and a defined set of CNA representations; support for more complex structural variant interpretation, including gene fusions and large rearrangements, falls outside the present scope. Finally, the use of a single prioritised transcript per variant is a deliberate trade-off between consistency and biological completeness (multi-transcript information remains accessible from the Tier 1 MAF) and the research-focused outputs are designed to support, rather than replace, expert scientific review by multidisciplinary research teams.

## Intended use statement

SMART is provided as a research tool for somatic variant annotation and exploratory analysis. It is not intended for, and must not be used for, patient-specific clinical reporting, treatment selection, diagnostic or prognostic decision-making, reimbursement, or regulatory decisions. Any clinical, patient-facing, or commercial use of SMART outputs, including outputs that contain or derive from OncoKB annotations, requires appropriate institutional validation and governance, professional review, and all required third-party licences, including any applicable OncoKB licence for generating patient reports.

## Acknowledgement

The authors would like to thank Wessex Genomics Laboratory Service for their support throughout the development of this work. The authors also acknowledge the use of the IRIDIS High Performance Computing Facility and associated support services at the University of Southampton in the completion of this work.

## Funding

This work is supported by funding from the Wessex Secure Data Environment and Central & South Genomic Medicine Service Alliance Research & Innovation.

## Conflict of Interest

The authors declare no competing interests.

## Data Availability Statement

The SMART source code, Docker image, reference installer scripts, configuration files, validation test suite, and the VCF files used to reproduce all validation analyses are freely available at https://github.com/WeTGI-colab/SMART. A version-tagged snapshot of the repository corresponding to this manuscript has been archived at Zenodo (DOI: 10.5281/zenodo.20445865). Detailed documentation of the verification framework, input data, expected outputs, and instructions to reproduce the validation results presented in this study is provided within the repository.

## Declaration of Generative AI Use

Generative AI tools were used during the preparation of this manuscript to enhance clarity and readability. All content produced with assistance from these tools was subsequently reviewed, revised, and verified by the authors, who assume full responsibility for the final manuscript.

## Supplementary information

Supplementary data are available online.

**Supplementary Figure S1 Column structure of the SMART Tier 1 MAF output.** Each block represents a functionally related group of annotation fields within the 1,028-column MAF file produced by SMART. Block colour denotes the originating annotation source or category. Column counts are shown in the top-right of each block. The OncoKB therapeutic section (TX 0–33) is the largest contributor, encoding 34 ranked treatment entries × 20 fields per entry (680 columns). MAF, Mutation Annotation Format; TFBS, transcription factor binding site; SV, structural variant; LoF, loss of function; AMP, Association for Molecular Pathology. A full description of all fields can be found on the SMART website (https://manuel-dominguezcbg.github.io/SMART/)

**Supplementary Figure S2 CPU and memory utilisation during the analysis of 100 VCF files comprising 254,662 variants.** Resource consumption was monitored throughout the execution of SMART using 8 parallel jobs on a desktop workstation (Intel Core i7-3770S, 8 cores, 3.10 GHz, 15 GB RAM). Peak memory usage remained below 11 GB during the analysis.

**Supplementary Figure S3. Comparative overview of PCGR, CancerVar, and SMART for somatic variant annotation and research interpretation.**

**Supplementary Table S1. Curated variant panel used for SMART pipeline verification.** All 22 variants are listed with their genomic coordinates (GRCh38/hg38), variant identifier, protein change, variant type, and predicted consequence. Gene names are reported using HGNC symbols. Protein changes follow HGVS nomenclature. CNA, copy-number alteration; SNV, single-nucleotide variant.

**Supplementary Table S2. VEP field-level concordance per variant.** For each of the 17 VEP-evaluable variants (5 excluded: 4 CNAs and GNAQ p.Gln209= owing to missing HGVSc), concordance is shown across 14 fields. P, concordant; S, skipped; U, untestable (INTRON field — no intronic variants in this panel).

**Supplementary Table S3. CIViC field-level concordance per variant.** CIViC was queried by protein change for the 18 SNV/indel variants (4 CNAs excluded). All 60 field checks passed (100% concordance). ERBB2 p.S310F was matched against CIViC variant id 497 (S310F/Y). P, concordant; S, skipped (CNA); —, no CIViC record in either MAF or API.

**Supplementary Table S4. OncoKB field-level concordance per variant.** All 22 variants were evaluated across 23 OncoKB fields. P, concordant; U, untestable (LEVEL_3B, ONCOKB_diagnosticSummary and ONCOKB_prognosticSummary are only populated when a specific tumour type is provided; all were consistently empty in both MAF and API under the generic query mode used here). The table continues on the following (landscape) page.

## Notes

https://zenodo.org/records/20445865

### Competing Interest Statement

The authors have declared no competing interest.

