## Supplementary Figure S1 for "SMART: A Somatic Mutation Annotation and Reporting Tool for cancer genomics"

### MAF core — 13 cols
Hugo symbol, sample barcode, coordinates, alleles, variant classification & type

### VCF raw — 8 cols
Preserved from input VCF: POS, ID, REF, ALT, QUAL, FILTER, FORMAT, FORMAT_DATA

### Genotype & sequencing metrics — 5 cols
Extracted from FORMAT field: AD (allele depth), DP (total depth), GT, PR, SR

### Structural variant fields — 5 cols
CIPOS, CIEND, END, SVLEN, SVTYPE - populated for SVs; empty for SNVs/indels

### VEP transcript annotation — 43 cols
Consequence, MANE Select, APPRIS, TSL, ENSP, UniProt, CCDS, REFSEQ_MATCH...

### VEP functional predictions — 5 cols
SIFT, PolyPhen-2 scores, protein DOMAINS, miRNA impact, HGVS_OFFSET flag

### VEP regulatory / motif — 9 cols
CLIN_SIG, SOMATIC, PHENO, PUBMED; TFBS motif name, position, score change

### Population allele frequencies — 29 cols
1000 Genomes (6 pops) · gnomAD exomes (10 pops) · gnomAD genomes (11 pops) · MAX_AF

### SpliceAI — 10 cols
Delta scores & positions for all four splice events: acceptor/donor gain & loss

### REVEL & LOEUF — 2 cols
REVEL ensemble missense pathogenicity · gnomAD LoF constraint metric

### ClinVar (detailed) — 40 cols
CLNSIG with SCVs, oncogenicity (ONC), somatic interpretation (SCI), review status...

### Cancer Hotspots — 10 cols
Recurrent SNV/indel hotspot · 3D structural hotspot · non-coding hotspot, with HGVSp/c

### CIViC summary & detailed — 41 cols
Top-level CIViC presence flag, gene name, and variant type and HGVS, evidence items, diseases, therapies, AMP category, ACMG code

### COSMIC — 8 cols
Variant/profile IDs and COSMIC annotation

### OncoKB core query & summary — 27 cols
Query params, hotspot, gene/variant/tumour summaries, highest Sens/Resist/FDA/Dx/Px levels

### OncoKB extended / mut. effect — 8 cols
Highest FDA level, other significant levels, mutation effect description + citations

### OncoKB diagnostic — 64 cols
4 ranked diagnostic entries × 16 fields: evidence level, tumour type (OncoTree), PMIDs

### OncoKB therapeutic (TX 0-33) - largest section — 680 cols
34 ranked treatment entries × 20 fields: drugs, approved indications, evidence level, FDA level, excluded cancer types, description, associated cancer type (OncoTree code, tissue, parent...)

### OncoKB flat summary levels — 27 cols
ANNOTATED, ONCOGENIC, MUTATION_EFFECT, LEVEL_1-4, R1-R2, Dx1-3, Px1-3, highest levels

Input / core · VEP annotation · Population frequencies · In silico predictors · Clinical databases · OncoKB
