## Supplementary figures and images for "SMART: A Somatic Mutation Annotation and Reporting Tool for cancer genomics"

### Supplementary Figure S2

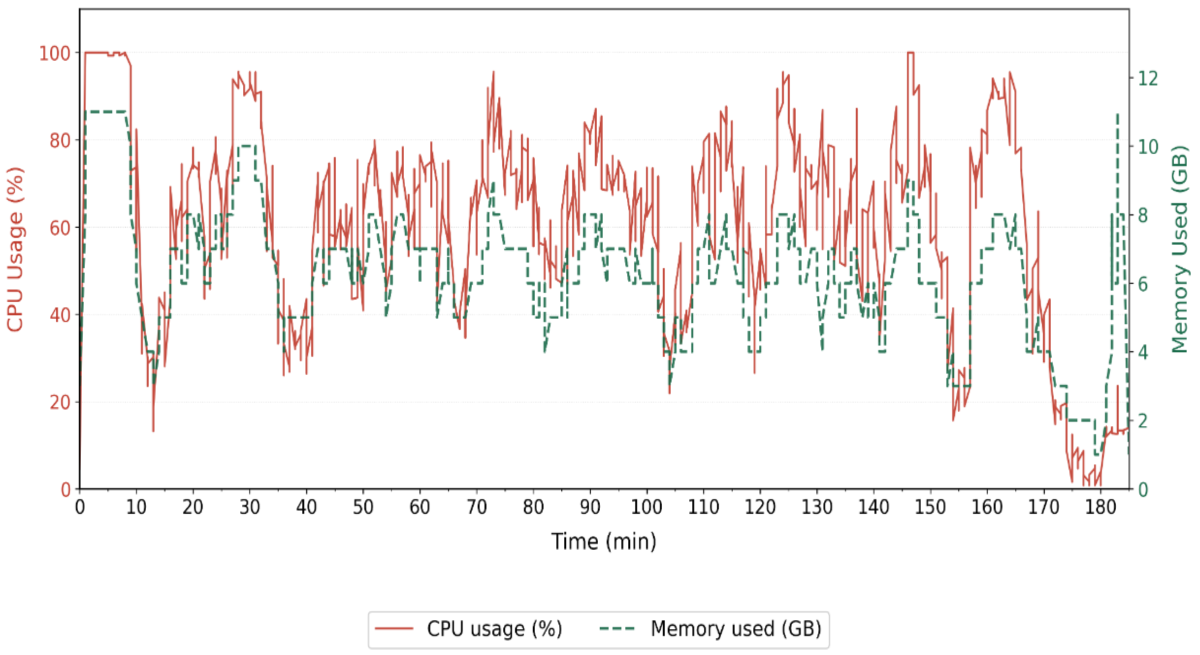
