## Supplementary Figure S3 for "SMART: A Somatic Mutation Annotation and Reporting Tool for cancer genomics"

| Category | Feature / Database | v2.2 | CancerVar v1.1 | SMART |
|---|---|---|---|---|
| **FUNCTIONAL ANNOTATION** | | | | |
| Functional consequence | VEP (variant effect prediction) | ✅ v113, GENCODE v47 | ✅ ANNOVAR | ✅ VEP v114, RefSeq / MANE |
| Functional consequence | Transcript prioritisation (MANE/whitelist) | MANE, MANE PLUS CLINICAL or Canonical | ❌ | ✅ Whitelist > MANE > VEP fallback |
| Protein domains | UniProt / SwissProt (2025_01) + Pfam (v37.0) | ✅ UniProt 2025_01 + Pfam v37.0 | ❌ | ✅ via VEP ((UniProt / InterPro) |
| **IN SILICO PREDICTIONS** | | | | |
| In silico predictions | dbNSFP (25 tools incl. CADD, AlphaMissense, ESM1b...) | ✅ dbNSFP v5.0 (25 scores) | ⚠️ dbnsfp30a/31a via ANNOVAR | ⚠️ SIFT & PolyPhen via VEP; REVEL separately |
| In silico predictions | REVEL (missense pathogenicity) | ⚠️ via dbNSFP v5.0 | ❌ | ✅ v1.3 (standalone VEP plugin) |
| In silico predictions | SpliceAI (splicing impact) | ⚠️ via dbNSFP v5.0 (splice_site_ADA/RF) | ❌ | ✅ v1.3 (standalone VEP plugin) |
| In silico predictions | CADD (combined annotation score) | ✅ via dbNSFP v5.0 | ✅ via ANNOVAR | ❌ |
| Gene constraint | LOEUF (gnomAD LoF intolerance) | ❌ | ❌ | ✅ gnomAD v4.0 (VEP plugin) |
| ML driver prediction | BoostDM / IntOGen (tumour-type driver scoring) | ❌ | ❌ | ❌ |
| **POPULATION FREQUENCIES** | | | | |
| Population frequencies | gnomAD | ✅ | ✅ gnomAD (via ANNOVAR) | ✅ v4.0 exomes + genomes |
| Population frequencies | dbSNP | ✅ build 156 | ✅ avsnp147 via ANNOVAR | ✅ via VEP |
| **SOMATIC VARIANT DATABASES** | | | | |
| Somatic hotspots | CancerHotspots (linear hotspots) | ✅ v2 (linear only) | ❌ | ✅ v2 linear (changv2_gao_nc) |
| Somatic hotspots | CancerHotspots (3-D structural hotspots) | ❌ | ❌ | ✅ 3D hotspots (Gao et al., Nat. Comms) |
| Tumour frequencies | TCGA (33 tumour types) | ✅ release 41.0 (Aug 2024) | ❌ | ❌ |
| Tumour frequencies | COSMIC | ⚠️ IDs via VEP Existing_variation | ✅ v91 via ANNOVAR | ✅ via COSMIC dataset v104 |
| Cancer driver genes | IntOGen / CancerMine | ✅ CancerMine v50 + IntOGen | ❌ | ❌ |
| **CLINICAL & THERAPEUTIC DATABASES** | | | | |
| Clinical significance | ClinVar | ✅ Mar 2025 | ✅ 2019-03-05 | ✅ Mar 2026 |
| Clinical evidence | CIViC | ✅ Mar 2025 | ❌ web server only | ✅ v3.6 |
| Clinical/therapeutic | CGI Cancer Biomarkers database | ✅ Oct 2022 | ❌ | ❌ |

| Category | Feature / Database | v2.2 | CancerVar v1.1 | SMART |
|---|---|---|---|---|
| Clinical/therapeutic | OncoKB | ❌ | ❌ external reference only (no native annotation) | ✅ (token required) |
| Oncogenes/suppressors | CancerMine | ✅ v50 | ❌ | ❌ |
| **VARIANT TYPES SUPPORTED** | | | | |
| Variant types | SNVs & small indels | ✅ | ✅ | ✅ |
| Variant types | Copy number alterations (CNV) | ✅ allele-specific segments | ✅ | ✅ Manta DUP / DEL |
| Variant types | Structural variants (BND, large INS) | ❌ | ❌ | ✅ MantaBND, MantaINS |
| Variant types | Gene fusions | ❌ | ⚠️ limited (interpretation only, not detection/annotation) | ❌ |
| Variant types | RNA-seq expression data | ✅ TCGA / DepMap / TreeHouse | ❌ | ❌ |
| **CLINICAL CLASSIFICATION FRAMEWORK** | | | | |
| Classification | Oncogenicity classification | ✅ VICC/ClinGen guidelines-informed | ✅ guideline-driven + ML (OPAI) | ✅ knowledgebase-driven via OncoKB |
| Therapeutic levels | Therapeutic sensitivity (Level 1–4) | ⚠️ via CIViC / CGI Does not strictly map to AMP levels 1–4, but provides evidence that can be interpreted into tiers | ⚠️ CBP[0] therapeutic evidence | ✅ OncoKB Level 1–4 |
| Therapeutic levels | Therapeutic resistance (Level R1–R3) | ⚠️ via CIViC Extracts resistance evidence from CIViC/CGI but does not standardise into R1–R3 tiers | ⚠️ CBP[0] therapeutic evidence but not standardised into AMP resistance tiers | ✅ OncoKB R1–R3 |
| Diagnostic/prognostic | Diagnostic implications (Dx tiers) | ❌ | ⚠️ CBP[1] diagnostic evidence Includes diagnostic evidence scoring, but not formally structured as Dx tiers | ✅ OncoKB Dx1–3 |
| Diagnostic/prognostic | Prognostic implications (Px tiers) | ❌ | ⚠️ CBP[2] prognostic evidence Not structured as Px tiers | ✅ OncoKB Px1–3 |
| Regulatory | FDA approval/companion diagnostic levels | ❌ | ❌ | ✅ OncoKB FDA levels |
| **TECHNICAL & DEPLOYMENT** | | | | |
| Input format | VCF input | ✅ | ✅ | ✅ |
| Coordinate handling | hg19 → hg38 liftover (built-in) | ⚠️ supports hg19/hg38 (no built-in liftover) | ⚠️ supports hg19/hg38 (no built-in liftover) | ✅ GATK LiftoverVcf v4.6 |

| Category | Feature / Database | v2.2 | CancerVar v1.1 | SMART |
| --- | --- | --- | --- | --- |
| Output format | Interactive HTML report | ✅ Quarto-based | ✅ Web interface | ❌ |
| Output format | TSV / MAF / VCF machine-readable output | ✅ TSV + VCF + Excel | ✅ TSV output (CancerVar-specific format) | ✅ MAF (Tier 1) TSV (Tier 2 and Tier 3) and VCF |
| Output format | Tiered output (clinician vs bioinformatician) | ❌ | ❌ | ✅ |
| Deployment | Docker / Singularity container | ✅ Docker + Singularity | ⚠️ Web + local install | ✅ Dockerised |
| Processing | Multi-sample batch processing | ❌ Per-sample | ❌ Per-sample | ✅ Per-sample or Cohort-level |
| Cancer types | Tumour-type aware annotation | ✅ 31 primary sites | ✅ ~30 cancer types (fixed categories) | ✅ 15 types |
