## Supplementary Data 1 for "SMART: A Somatic Mutation Annotation and Reporting Tool for cancer genomics"

Field-level verification of the SMART annotation pipeline against reference APIs

This Supplementary Data file contains the complete variant-level and field-level concordance results from the SMART pipeline verification study. Verification was performed by querying the OncoKB, VEP, and CIViC reference APIs directly for each variant in the curated panel and comparing the returned values against those stored in the pipeline-generated MAF file. A total of 804 field checks were performed across all three modules, with 100% concordance and zero mismatches or coverage gaps. Status codes used throughout are: P, concordant (PASS); S, skipped (variant type not evaluable by the given module, e.g. copy-number alterations for VEP and CIViC); U, untestable (field always empty in both MAF and API for this dataset); —, no record present in either MAF or API (consistent absence, counted as PASS).

**Supplementary Table S1 |** Curated variant panel used for SMART pipeline verification.

All 22 variants are listed with their genomic coordinates (GRCh38/hg38), variant identifier, protein change, variant type, and predicted consequence. Gene names are reported using HGNC symbols. Protein changes follow HGVS nomenclature. CNA, copy-number alteration; SNV, single-nucleotide variant.

| **#** | **Gene** | **Variant ID** | **Protein change** | **Variant type** | **Consequence** | **Chromosome: position** |
| --- | --- | --- | --- | --- | --- | --- |
| 1 | *NRAS* | NRAS_Q61R | p.Q61R | SNV | Missense | chr1:114713908 T>C |
| 2 | *IDH1* | IDH1_R132H | p.R132H | SNV | Missense | chr2:208248388 C>T |
| 3 | *PIK3CA* | PIK3CA_E545K | p.E545K | SNV | Missense | chr3:179218303 G>A |
| 4 | *PIK3CA* | PIK3CA_H1047R | p.H1047R | SNV | Missense | chr3:179234297 A>G |
| 5 | *EGFR* | EGFR_del19 | p.Glu746_Ala750del | Indel | In-frame deletion | chr7:55174772 |
| 6 | *EGFR* | EGFR_T790M | p.T790M | SNV | Missense | chr7:55181378 C>T |
| 7 | *EGFR* | EGFR_L858R | p.L858R | SNV | Missense | chr7:55191822 T>G |
| 8 | *MET* | MantaDUP: MET_AMP | — | CNA | Amplification | chr7:116672196 |
| 9 | *BRAF* | BRAF_V600E | p.V600E | SNV | Missense | chr7:140753336 A>T |
| 10 | *CDKN2A* | MantaDEL:CDKN2A_DEL | — | CNA | Deletion | chr9:21967753 |
| 11 | *GNAQ* | GNAQ_Q209L | p.Gln209= | SNV | Synonymous | chr9:77794572 A>T |
| 12 | *PTEN* | PTEN_R130stop | p.R130* | SNV | Stop gained | chr10:87933147 C>T |
| 13 | *KRAS* | KRAS_G13D | p.G13D | SNV | Missense | chr12:25245347 C>T |
| 14 | *KRAS* | KRAS_G12D | p.G12D | SNV | Missense | chr12:25245350 C>T |
| 15 | *KRAS* | KRAS_G12C | p.G12C | SNV | Missense | chr12:25245351 C>A |
| 16 | *CDK4* | MantaDUP:CDK4_AMP | — | CNA | Amplification | chr12:57748512 |
| 17 | *BRCA2* | BRCA2_L3101R | p.L3101R | SNV | Missense | chr13:32394734 T>G |
| 18 | *DICER1* | DICER1_E1705K | p.Glu1705* | SNV | Stop gained | chr14:95094139 G>A |
| 19 | *TP53* | TP53_R248W | p.R248W | SNV | Missense | chr17:7674221 G>A |
| 20 | *TP53* | TP53_R175H | p.R175H | SNV | Missense | chr17:7675088 C>T |
| 21 | *ERBB2* | MantaDUP:ERBB2_AMP | — | CNA | Amplification | chr17:39687914 |
| 22 | *ERBB2* | ERBB2_S310F | p.S310F | SNV | Missense | chr17:39711955 C>T |

**Supplementary Table S2 |** VEP field-level concordance per variant.

For each of the 17 VEP-evaluable variants (5 excluded: 4 CNAs and GNAQ p.Gln209= owing to missing HGVSc), concordance is shown across 14 fields. P, concordant; S, skipped; U, untestable (INTRON field — no intronic variants in this panel).

| **Variant** | **Consequence** | **IMPACT** | **BIOTYPE** | **EXON** | **INTRON** | **HGVSc** | **HGVSp** | **Protein_position** | **Amino_acids** | **Codons** | **SIFT** | **PolyPhen** | **STRAND** | **CANONICAL** |
| --- | --- | --- | --- | --- | --- | --- | --- | --- | --- | --- | --- | --- | --- | --- |
| *NRAS Q61R* | P | P | P | P | U | P | P | P | P | P | P | P | P | P |
| *IDH1 R132H* | P | P | P | P | U | P | P | P | P | P | P | P | P | P |
| *PIK3CA E545K* | P | P | P | P | U | P | P | P | P | P | P | P | P | P |
| *PIK3CA H1047R* | P | P | P | P | U | P | P | P | P | P | P | P | P | P |
| *EGFR del19* | P | P | P | P | U | P | P | P | P | P | P | P | P | P |
| *EGFR T790M* | P | P | P | P | U | P | P | P | P | P | P | P | P | P |
| *EGFR L858R* | P | P | P | P | U | P | P | P | P | P | P | P | P | P |
| *MET AMP* | S | S | S | S | S | S | S | S | S | S | S | S | S | S |
| *BRAF V600E* | P | P | P | P | U | P | P | P | P | P | P | P | P | P |
| *CDKN2A DEL* | S | S | S | S | S | S | S | S | S | S | S | S | S | S |
| *GNAQ Q209L* | S | S | S | S | S | S | S | S | S | S | S | S | S | S |
| *PTEN R130** | P | P | P | P | U | P | P | P | P | P | P | P | P | P |
| *KRAS G13D* | P | P | P | P | U | P | P | P | P | P | P | P | P | P |
| *KRAS G12D* | P | P | P | P | U | P | P | P | P | P | P | P | P | P |
| *KRAS G12C* | P | P | P | P | U | P | P | P | P | P | P | P | P | P |
| *CDK4 AMP* | S | S | S | S | S | S | S | S | S | S | S | S | S | S |
| *BRCA2 L3101R* | P | P | P | P | U | P | P | P | P | P | P | P | P | P |
| *DICER1 E1705** | P | P | P | P | U | P | P | P | P | P | P | P | P | P |
| *TP53 R248W* | P | P | P | P | U | P | P | P | P | P | P | P | P | P |
| *TP53 R175H* | P | P | P | P | U | P | P | P | P | P | P | P | P | P |
| *ERBB2 AMP* | S | S | S | S | S | S | S | S | S | S | S | S | S | S |
| *ERBB2 S310F* | P | P | P | P | U | P | P | P | P | P | P | P | P | P |

**Supplementary Table S3 |** CIViC field-level concordance per variant.

CIViC was queried by protein change for the 18 SNV/indel variants (4 CNAs excluded). All 60 field checks passed (100% concordance). ERBB2 p.S310F was matched against CIViC variant id 497 (S310F/Y). P, concordant; S, skipped (CNA); —, no CIViC record in either MAF or API.

| **Variant** | **CIViC Variant Name** | **SYMBOL** | **CIViC Variant ID** | **CIViC Molecular Profile Name** |
| --- | --- | --- | --- | --- |
| *NRAS Q61R* | P | P | P | P |
| *IDH1 R132H* | P | P | P | P |
| *PIK3CA E545K* | P | P | P | P |
| *PIK3CA H1047R* | P | P | P | P |
| *EGFR del19* | — | — | — | — |
| *EGFR T790M* | P | P | P | P |
| *EGFR L858R* | P | P | P | P |
| *MET AMP* | S | S | S | S |
| *BRAF V600E* | P | P | P | P |
| *CDKN2A DEL* | S | S | S | S |
| *GNAQ Q209L* | — | — | — | — |
| *PTEN R130** | P | P | P | P |
| *KRAS G13D* | P | P | P | P |
| *KRAS G12D* | P | P | P | P |
| *KRAS G12C* | P | P | P | P |
| *CDK4 AMP* | S | S | S | S |
| *BRCA2 L3101R* | — | — | — | — |
| *DICER1 E1705** | — | — | — | — |
| *TP53 R248W* | P | P | P | P |
| *TP53 R175H* | P | P | P | P |
| *ERBB2 AMP* | S | S | S | S |
| *ERBB2 S310F* | P | P | P | P |

**Supplementary Table S4 |** OncoKB field-level concordance per variant.

All 22 variants were evaluated across 23 OncoKB fields. P, concordant; U, untestable (LEVEL_3B, ONCOKB_diagnosticSummary and ONCOKB_prognosticSummary are only populated when a specific tumour type is provided; all were consistently empty in both MAF and API under the generic query mode used here). The table continues on the following (landscape) page.

| **Variant** | **ALLELE_EXIST** | **GENE_SUMMARY** | **HOTSPOT** | **VUS** | **highestFdaLevel** | **VARIANT_SUMMARY** | **GENE_IN_ONCOKB** | **VARIANT_IN_ONCOKB** | **MUTATION_EFFECT** | **ONCOGENIC** | **LEVEL_1** | **LEVEL_2** | **LEVEL_3A** | **LEVEL_3B** | **LEVEL_4** | **LEVEL_R1** | **LEVEL_R2** | **H_SENSITIVE_LVL** | **H_RESISTANCE_LVL** | **DIAG_LVL** | **PROG_LVL** | **Diagnostic Summary** | **Prognostic Summary** |
| --- | --- | --- | --- | --- | --- | --- | --- | --- | --- | --- | --- | --- | --- | --- | --- | --- | --- | --- | --- | --- | --- | --- | --- |
| *NRAS Q61R* | P | P | P | P | P | P | P | P | P | P | P | P | P | U | P | P | P | P | P | P | P | U | U |
| *IDH1 R132H* | P | P | P | P | P | P | P | P | P | P | P | P | P | U | P | P | P | P | P | P | P | U | U |
| *PIK3CA E545K* | P | P | P | P | P | P | P | P | P | P | P | P | P | U | P | P | P | P | P | P | P | U | U |
| *PIK3CA H1047R* | P | P | P | P | P | P | P | P | P | P | P | P | P | U | P | P | P | P | P | P | P | U | U |
| *EGFR del19* | P | P | P | P | P | P | P | P | P | P | P | P | P | U | P | P | P | P | P | P | P | U | U |
| *EGFR T790M* | P | P | P | P | P | P | P | P | P | P | P | P | P | U | P | P | P | P | P | P | P | U | U |
| *EGFR L858R* | P | P | P | P | P | P | P | P | P | P | P | P | P | U | P | P | P | P | P | P | P | U | U |
| *MET AMP* | P | P | P | P | P | P | P | P | P | P | P | P | P | U | P | P | P | P | P | P | P | U | U |
| *BRAF V600E* | P | P | P | P | P | P | P | P | P | P | P | P | P | U | P | P | P | P | P | P | P | U | U |
| *CDKN2A DEL* | P | P | P | P | P | P | P | P | P | P | P | P | P | U | P | P | P | P | P | P | P | U | U |
| *GNAQ Q209L* | P | P | P | P | P | P | P | P | P | P | P | P | P | U | P | P | P | P | P | P | P | U | U |
| *PTEN R130** | P | P | P | P | P | P | P | P | P | P | P | P | P | U | P | P | P | P | P | P | P | U | U |
| *KRAS G13D* | P | P | P | P | P | P | P | P | P | P | P | P | P | U | P | P | P | P | P | P | P | U | U |
| *KRAS G12D* | P | P | P | P | P | P | P | P | P | P | P | P | P | U | P | P | P | P | P | P | P | U | U |
| *KRAS G12C* | P | P | P | P | P | P | P | P | P | P | P | P | P | U | P | P | P | P | P | P | P | U | U |
| *CDK4 AMP* | P | P | P | P | P | P | P | P | P | P | P | P | P | U | P | P | P | P | P | P | P | U | U |
| *BRCA2 L3101R* | P | P | P | P | P | P | P | P | P | P | P | P | P | U | P | P | P | P | P | P | P | U | U |
| *DICER1 E1705** | P | P | P | P | P | P | P | P | P | P | P | P | P | U | P | P | P | P | P | P | P | U | U |
| *TP53 R248W* | P | P | P | P | P | P | P | P | P | P | P | P | P | U | P | P | P | P | P | P | P | U | U |
| *TP53 R175H* | P | P | P | P | P | P | P | P | P | P | P | P | P | U | P | P | P | P | P | P | P | U | U |
| *ERBB2 AMP* | P | P | P | P | P | P | P | P | P | P | P | P | P | U | P | P | P | P | P | P | P | U | U |
| *ERBB2 S310F* | P | P | P | P | P | P | P | P | P | P | P | P | P | U | P | P | P | P | P | P | P | U | U |
